# *De novo* assembly of the Mongolian gerbil genome and transcriptome

**DOI:** 10.1101/522516

**Authors:** Shifeng Cheng, Yuan Fu, Yaolei Zhang, Wenfei Xian, Hongli Wang, Benedikt Grothe, Xin Liu, Xun Xu, Achim Klug, Elizabeth A McCullagh

## Abstract

**BACKGROUND:** The Mongolian gerbil (*Meriones unguiculatus*) has historically been used as a model organism for the auditory and visual systems, stroke/ischemia, epilepsy and aging related research since 1935 when laboratory gerbils were separated from their wild counterparts. In this study we report genome sequencing, assembly, and annotation further supported by transcriptome data from 27 different tissues samples.

**FINDINGS:** The genome was assembled using Illumina HiSeq 2000 and resulted in a final genome size of 2.54 Gbp with contig and scaffold N50 values of 31.4 Kbp and 500.0 Kbp, respectively. Based on the k-mer estimated genome size of 2.48 Gbp, the assembly appears to be complete. The genome annotation was supported by transcriptome data that identified 36 019 predicted protein-coding genes across 27 tissue samples. A BUSCO search of 3023 mammalian groups resulted in 86% of curated single copy orthologs present among predicted genes, indicating a high level of completeness of the genome.

**CONCLUSIONS:** We report a *de novo* assembly of the Mongolian gerbil genome that was further enhanced by annotation of transcriptome data from several tissues. Sequencing of this genome increases the utility of the gerbil as a model organism, opening the availability of now widely used genetic tools.

The data sets supporting the results of this article are available in the China National GeneBank CNSA repository, Accession id: CNP0000340.

## DATA DESCRIPTION

### Background information on *Meriones unguiculatus*

The Mongolian gerbil is a small rodent that is native to Mongolia, southern Russia, and northern China. Laboratory gerbils used as model organisms originated from 20 founders captured in Mongolia in 1935 [1]. Gerbils have been used as model organisms for sensory systems (visual and auditory) and pathologies (aging, epilepsy, irritable bowel syndrome and stroke/ischemia). The gerbil’s hearing range covers the human audiogram while also extending into ultrasonic frequencies, making gerbils a better model than rats or mice to study lower frequency human-like hearing [2]. In addition to the auditory system, the gerbil has also been used as a model for the visual system because gerbils are diurnal and therefore have more cone receptors than mice or rats making them a closer model to the human visual system [3]. The gerbil has also been used as a model for aging due to its ease of handling, prevalence of tumors, and experimental stroke manipulability [1,4]. Interestingly, the gerbil has been used as a model for stroke and ischemia due to variations in the blood supply to the brain due to an anatomical region known as the “Circle of Willis” [5]. In addition, the gerbil is a model for epileptic activity as a result of its natural minor and major seizure propensity when exposed to novel stimuli [6,7]. Lastly, the gerbil has been used as model for inflammatory bowel disease, colitis, and gastritis due to the similarity in the pathology of these diseases between humans and gerbils [8,9]. Despite its usefulness as a model for all these systems and medical conditions, the utility of the gerbil as a model organism has been limited due to a lack of a sequenced genome to manipulate. This is especially the case with the increased use of genetic tools to manipulate model organisms.

Here we describe a *de novo* assembly and annotation of the Mongolian gerbil genome and transcriptome. Recently, a separate group has sequenced the gerbil genome, however our work is further supported by comparisons with an in-depth transcriptome analysis [10]. RNA-seq data were produced from 27 tissues that were used in the genome annotation and deposited in the NCBI SRA database under the project____. These data provide a draft genome sequence to facilitate the continued use of the Mongolian gerbil as a model organism and to help broaden the genetic rodent models available to researchers.

### Animals and Genome Sequencing

All experiments complied with all applicable laws, NIH guidelines, and were approved by the University of Colorado IACUC. Five young adult (postnatal day 65-71) gerbils (three males and two females) were used for tissue RNA transcriptome analysis and DNA genome assembly. In addition, two old (postnatal day 1013 or 2.7 years) female gerbil’s tissue was used for transcriptome analysis.

Genomic DNA was extracted from young adult animal tail and ear snips using a commercial kit (DNeasy Blood and Tissue Kit, Qiagen, Venlo, Netherlands). We then used the extracted DNA to create different pair-end insert libraries of 250 bp, 350 bp, 500 bp, 800 bp, 2 Kb, 4 Kb, 6 Kb, and 10 Kb. These libraries were then sequenced using an Illumina HiSeq2000 Genome Analyzer (Ilumina, San Diego, CA, USA) generating a total of 322.13 Gb in raw data, from which a total of 287.4 Gb of ‘clean’ data was obtained after removal of duplicates, contaminated reads, and low-quality reads.

### Assembly

The gerbil genome was estimated to be approximately 2.48 Gbp using a k-mer-based approach. High-quality reads were then used for genome assembly using the SOAPdenovo (version 2.04) package. The final assembly had a total length of 2.54 Gb and was comprised of 31,769 scaffolds assembled from 114,522 contigs. The N50 sizes for contigs and scaffolds were 31.4 Kbp and 500.0 Kbp, respectively (Table 1). Given the genome size estimate of 2.48 Gbp, genome coverage by the final assembly was likely complete and is consistent with the previously published gerbil genome, which had a total length of 2.523 Gbp [10]. Completeness of the genome assembly was confirmed by successful mapping of the RNA-seq assembly back to the genome showing that 98% of the RNA-seq sequences can be mapped to the genome with >50% sequence in one scaffold.

**Table 1:** Global statistics of the Mongolian gerbil genome

| Statistic | Value |
| --- | --- |
| Size (Gb) | 2.54 |
| Scaffold number (>2000bp) | 31769 |
| Scaffold N50 (Kb) | 500.0 |
| Contig number (>2000bp) | 114522 |
| Contig N50 (Kb) | 31.4 |

### Transcriptome Sequencing/Assembly/Annotation

Gene expression data were produced to aid in the genome annotation process. Samples from 27 tissues were collected from the seven gerbils described above (Supplementary Figure 1). The tissues were collected after the animals were euthanized with isoflurane and stored on liquid nitrogen until homogenized with a pestle. RNA was prepared using the RNeasy mini isolation kit (Qiagen, Venlo, Netherlands). RNA integrity was analyzed using a Nanodrop Spectrophotometer (Thermo Fisher Waltham, MA, USA) followed by analysis with an Agilent Technologies 2100 Bioanalyzer (Agilent Technologies, Santa Clara, CA, USA) and samples with an RNA integrity number (RIN) value greater than 7.0 were used to prepare libraries which were sequenced using an Ilumina Hiseq2000 Genome Analyzer (Ilumina, San Diego, CA, USA). The sequenced libraries were assembled with Trinity (v2.0.6 parameters: "--min_contig_length 150 --min_kmer_cov 3 --min_glue 3 --bfly_opts '-V 5 --edge-thr=0.1 --stderr'") generating 131,845 sequences with a total length of 130,734,893 bp. Quality of the RNA assembly was assessed by filtering RNA-seq reads using SOAPnuke (v1.5.2 parameters: "-l 10 -q 0.1 -p 50 -n 0.05 -t 5,5,5,5") followed by mapping of clean reads to the assembled genome using HISAT2 (v2.0.4) and StringTie (v1.3.0). The initial assembled genes were then filtered using CD-HIT (v4.6.1) with sequence identity threshold of 0.9 followed by a homology search (human, rat, mouse proteins) and TransDecoder (v2.0.1) open reading frame (ORF) prediction. The RNA-seq assembly resulted in 19,737 protein-coding genes with a total length of 29.4 Mbp, which is available in the China National GeneBank CNSA repository, Accession id: CNP0000340.

**Figure 1:**
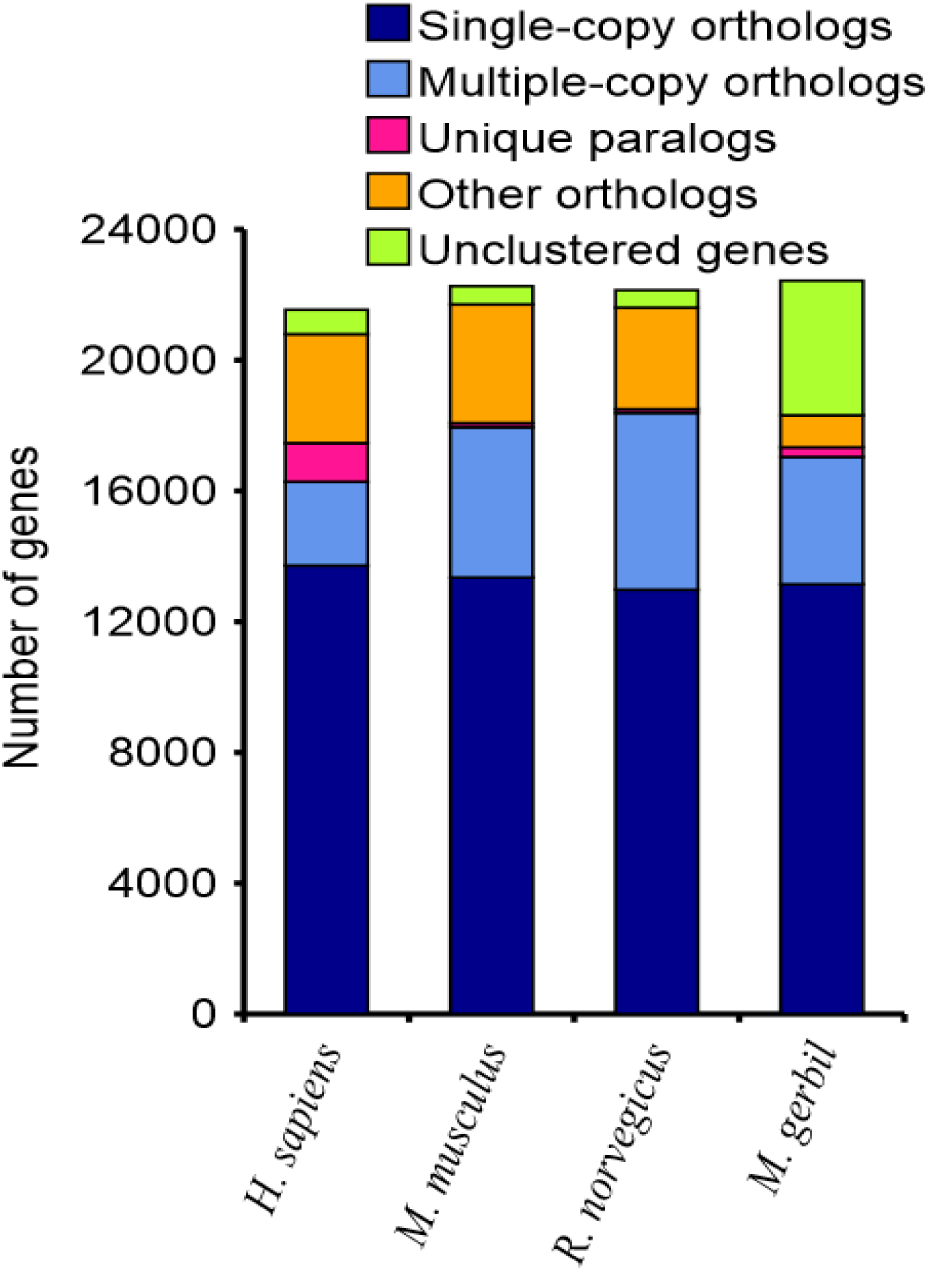
Gene Family Construction. The number of genes is similar between species compared (human, mouse, rat, and gerbil.

### Genome Annotation

Genomic repeat elements of the genome assembly were also identified and annotated using RepeatMasker (v4.0.5 RRID:SCR_012954)[11] and RepBase library (v20.04)[12]. In addition, we constructed a *de novo* repeat sequence database using LTR-FINDER (v1.0.6) [13] and RepeatModeler (v1.0.8) [13] to identify any additional repeat elements using RepeatMasker. A combination of both repeat element identification approaches resulted in a total length of 1016.7 Mbp of the total *M. unguiculatus* genome as repetitive, accounting for 40.0% of the entire genome assembly. The repeat element landscape of *M. unguiculatus* consists of long interspersed elements (LINEs)(27.5%), short interspersed elements (SINEs)(3.7%), long terminal repeats (LTRs)(6.5%), and DNA transposons (0.81%) (Table 2). This is consistent with other rodent species including mouse [14] and rat [15].

**Table 2:**
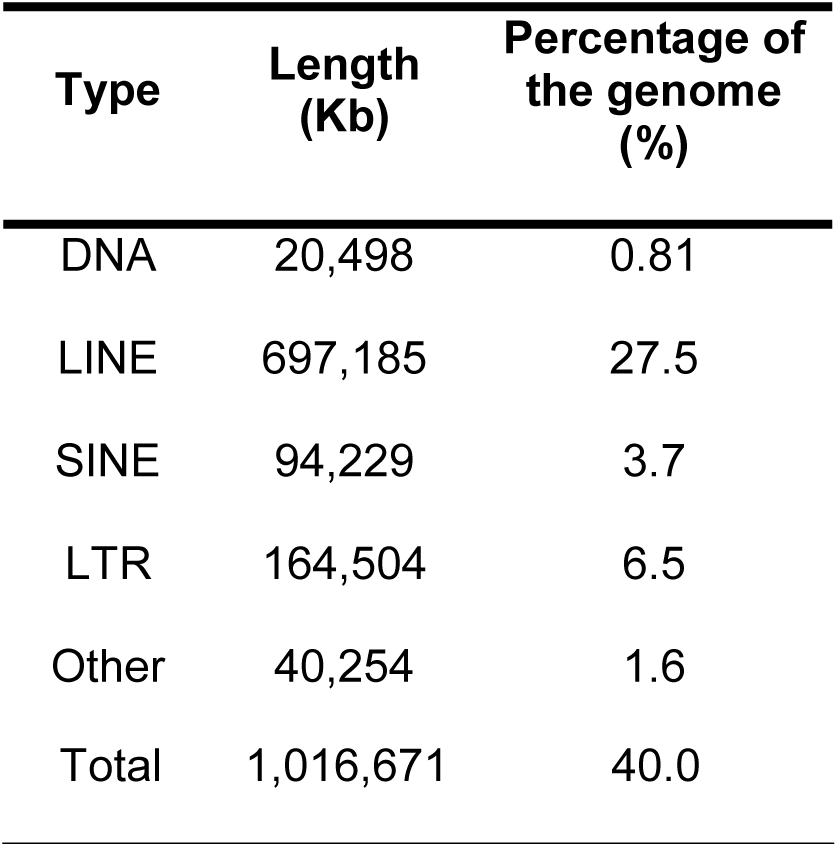
Summary of mobile element types

Protein-coding genes were predicted and annotated by a combination of homology searching, *ab initio* prediction (using AUGUSTUS (v3.1), GENSCAN (1.0), and SNAP (v2.0)), and RNA-seq data (using TopHat (v1.2 with parameters: “-p 4 --max-intron-length 50000 -m 1 –r 20 --mate-std-dev 20 --closure-search --coverage-search --microexon-search”) and Cufflinks (v2.2.1 http://cole-trapnell-lab.github.io/cufflinks/)) after repetitive sequences in the genome were masked using known repeat information detected by RepeatMasker and RepeatProteinMask. Homology searching was performed using protein data from *Homo Sapiens* (human), *Mus musculus* (mouse), *and Rattus norvegicus* (rat) from Ensembl (v80) aligned to the masked genome using BLAT. Genewise (v2.2.0) was then used to improve the accuracy of alignments and to predict gene models. The *de novo* gene predictions and homology-based search were then combined using GLEAN. The GLEAN results were then integrated with the transcriptome dataset using an in-house program (Table 3). This resulted in an identification of a total of 22,998 protein-coding genes with an average transcript length of 23,846.58 bp. There were an average of 7.76 exons per gene with an average length of 197.9 bp and average intron length of 3300.83 bp. The 22,998 protein-coding genes were aligned to several protein databases to begin to identify their possible function. InterProScan (v5.11) was used to align the final gene models to databases (ProDom, ProSiteProfiles, SMART, PANTHER, PRINTS, Pfam, PIRSF, ProSitePatterns, SignalP_EUK, Phobius, IGRFAM, and TMHMM) to detect consensus motifs and domains within these genes. Using the InterProScan results, we obtained the annotations of the gene products from the Gene Ontology database. We then mapped these genes to proteins in SwissProt and TrEMBL (Uniprot release 2015.04) using blastp with an E-value <1E-5. We also aligned the final gene models to proteins in KEGG (release 76) to determine the functional pathways for each gene (Table 4). This resulted in 20,760 protein-coding genes that had a functional annotation, or 90.3% of the total gene set.

**Table 3:** General statistics of predicted protein-coding genes

|  | Gene set | Number | Average transcript length (bp) | Average CDS length (bp) | Average exon per gene | Average exon length (bp) | Average intron length (bp) |
| --- | --- | --- | --- | --- | --- | --- | --- |
| De novo | SNAP | 76858 | 42227.63 | 742.83 | 5.52 | 134.62 | 9182.18 |
|  | AUGUSTUS | 24675 | 19838.68 | 1133.22 | 5.61 | 201.97 | 4056.79 |
|  | GENESCAN | 49390 | 24183.55 | 1023.1 | 6.25 | 163.54 | 4406.54 |
| Homolog | *Mus musculus* | 22728 | 26977.32 | 1465.18 | 8.02 | 182.61 | 3632.46 |
|  | *Rattus norvegicus* | 23686 | 23564.96 | 1336.56 | 7.43 | 179.83 | 3455.8 |
|  | *Homo sapiens* | 17131 | 31217.18 | 1580.27 | 9.11 | 173.55 | 3656.27 |
|  | GLEAN | 19893 | 18835.39 | 1418.26 | 7.72 | 183.69 | 2691.49 |
|  | Transcriptome | 36019 | 33752.29 | 1758.58 | 10.74 | 163.77 | 3285.43 |
|  | Final set | 22998 | 23846.58 | 1535.48 | 7.76 | 197.9 | 3300.83 |

**Table 4:**
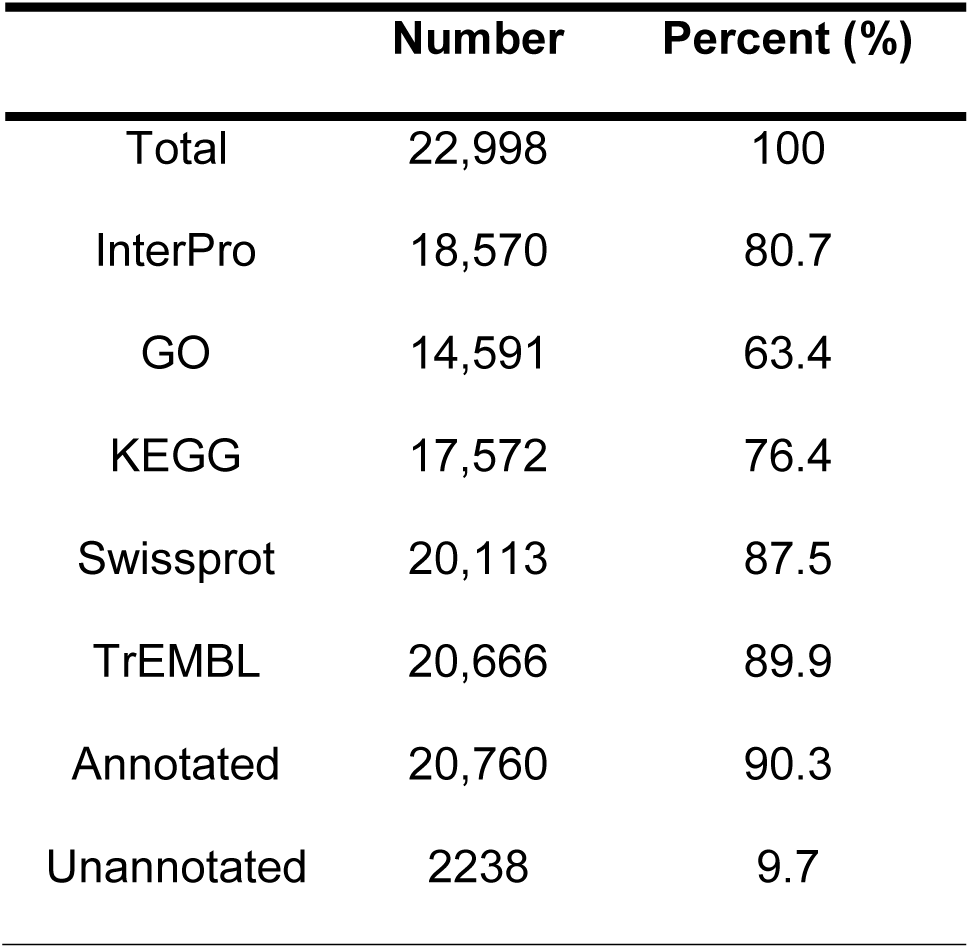
Functional annotation of the final gene set

| Number | Percent (%) |
| --- | --- |

|  |  |  |
| --- | --- | --- |
| Total | 22,998 | 100 |
| InterPro | 18,570 | 80.7 |
| GO | 14,591 | 63.4 |
| KEGG | 17,572 | 76.4 |
| Swissprot | 20,113 | 87.5 |
| TrEMBL | 20,666 | 89.9 |
| Annotated | 20,760 | 90.3 |
| Unannotated | 2238 | 9.7 |

### Quality Assessment

In addition to measuring standard assembly quality metrics, genome assembly and annotation quality were further assessed by comparison with closely related species, gene family construction, evaluation of housekeeping genes, and Benchmarking Universal Single-Copy Orthologs (BUSCO) search. The assembled gerbil genome was compared with other closely related model organisms including mouse, rat, and hamster (Table 5). The genomes from these species varied in size from 2.3 to 2.8 Gbp. The total number of annotated proteins in gerbil (20,760) is most similar to mouse (22,598), followed by rat (23,347), and then hamster (24,238). Gene family construction was performed using Treefam (http://www.treefam.org/) (Figure 1). This analysis showed that single-copy orthologs in gerbil are similar to mouse and rat. To examine housekeeping genes we downloaded 2169 human housekeeping genes from (http://www.tau.ac.il/~elieis/HKG/) and extracted corresponding protein sequences to align to the gerbil genome using blastp (v.2.2.26). We found there were 2141 genes consistent between human and gerbil housekeeping genes (this is similar to rat (2153) and mouse (2146). Lastly, we employed BUSCO (v1.2) to search 3023 mammalian groups. Of these groups, 86% complete BUSCO groups can be detected in the final gene set. The presence of 86% complete mammalian BUSCO gene groups suggests a high level of completeness of this gerbil genome assembly. A BUSCO search was also performed for the gerbil transcriptome data resulting in detection of 82% complete BUSCO groups in the final transcriptome dataset (Table 6). Based on the results from the quality metrics described above, we are confident of the quality of the data for this assembly of the gerbil genome and transcriptome.

**Table 5:** Genome annotation comparisons with other model organisms

| Species | Common name | Protein coding genes | Assembly Size | Divergence time to gerbils, Myr | RefSeq assembly accession | Annotation release ID |
| --- | --- | --- | --- | --- | --- | --- |
| *Meriones unguiculatus* | Mongolian gerbil | 20,760 | 2,537,533,819 | -- |  |  |
| *Mus musculus* | mouse | 22,598 | 2,818,974,548 | 22.5 | GCF_000001635.26 | 106 |
| *Rattus norvegicus* | rat | 23,347 | 2,870,184,193 | 22.5 | GCF_000001895.5 | 106 |
| *Cricetulus griseus* | Chinese hamster | 24,238 | 2,358,151,106 | 25 | GCA_900186095.1 | 102 |

**Table 6:** Completeness of gerbil genome and transcriptome assembly as assessed by BUSCO

|  | Genome | Transcriptome |
| --- | --- | --- |
| Complete BUSCOs | 2601 | 2508 |
| Duplicated BUSCOs | 55 | 46 |
| Fragmented BUSCOs | 170 | 293 |
| Missing BUSCOs | 252 | 222 |
| Total BUSCO groups searched | 3023 | 3023 |

In summary, we report a fully annotated Mongolian gerbil genome sequence assembly enhanced by transcriptome data from several different gerbils and tissues. The gerbil genome and transcriptome adds to the availability of alternative rodent models that may be better models for diseases than rats or mice. Additionally, the gerbil is an interesting comparative rodent model to mouse and rat since it has many traits in common, but also differs in seizure susceptibility, low-frequency hearing, cone visual processing, stroke/ischemia susceptibility, gut disorders and aging. Sequencing of the gerbil genome and transcriptome opens these areas to molecular manipulation in the gerbil and therefore better models for specific disease states.

### Availability of supporting data

Genome annotation results are available at the China National GeneBank CNSA repository, Accession id: CNP0000340, and supporting materials, which include transcripts and genome assembly, are available under the same project.

## Additional files

Additional file 1: Table S1 Tissues analyzed for RNA-seq data

## Abbreviations

bp: base pair
BUSCO: Benchmarking Universal Single-Copy Orthologs
CDS: coding sequence
LINEs: long interspersed elements
LTRs: long terminal repeats
Myr: million years
NCBI: National Center for Biotechnology Information
RefSeq: Reference sequence
RNA-seq: high-throughput messenger RNA sequencing
RIN: RNA integrity number
SINEs: short interspersed elements

## Authors’ contributions

SC, EAM, and AK developed the ideas, methods, and, wrote and revised the manuscript. BG, YF, YZ, WX, HW, XL, and XX advised and revised the manuscript. BG provided the old animal tissues from Munich, Germany. SC, YF, YZ, WX, HW, XL, and XX performed the analysis and annotation of the genome and transcriptome. EAM prepared the DNA and RNA samples for sequencing.

## Acknowledgements

The authors would like to thank Hilde Wohlfrom for sending tissues from Germany.

---

**Figure S1:**
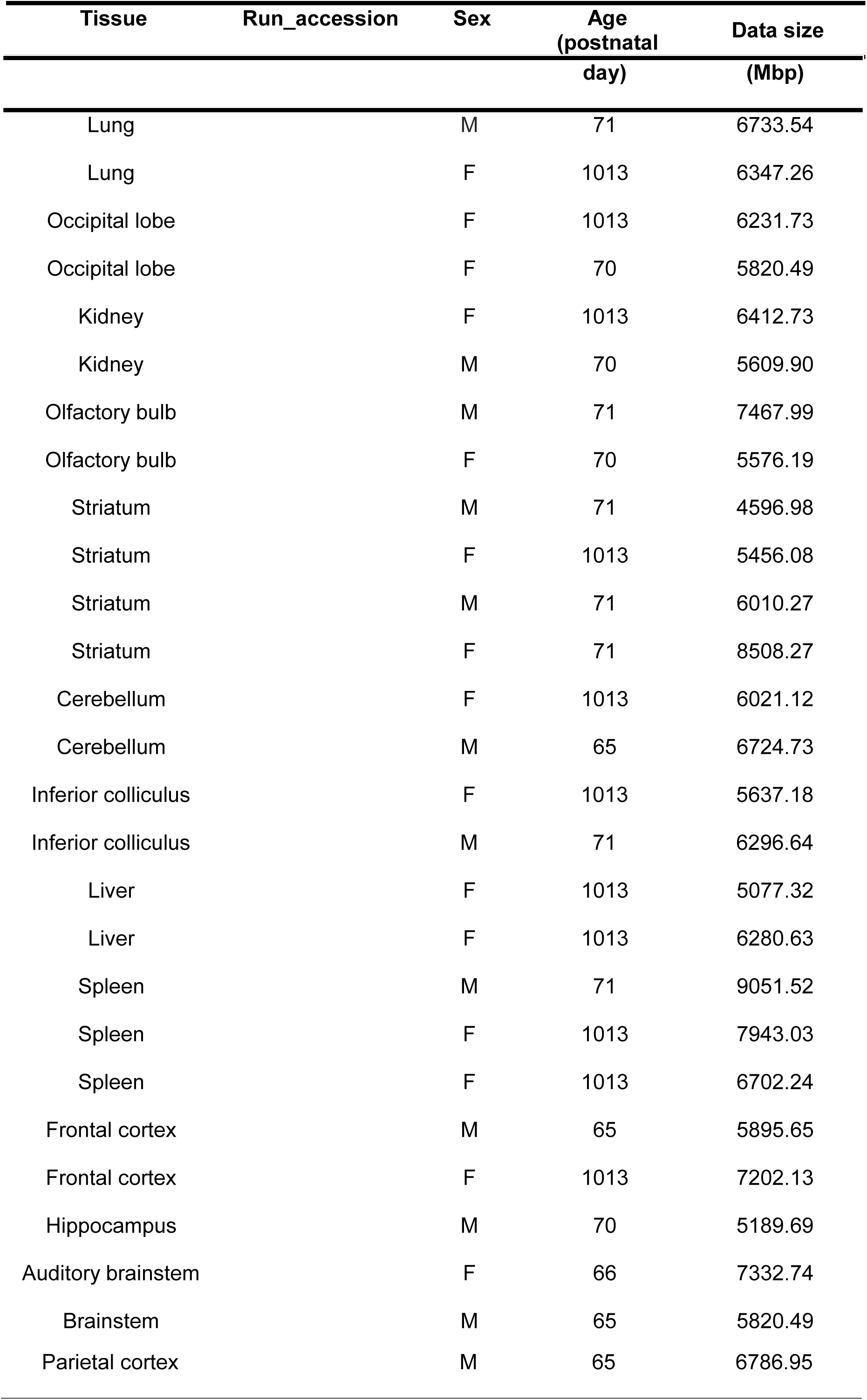
Tissues sampled for RNA transcriptome

